# Conventional dendritic cells 2, the targeted antigen-presenting-cell, induces enhanced type 1 immune responses in mice immunized with CVC1302 in oil formulation

**DOI:** 10.1101/2022.10.24.513633

**Authors:** Luping Du, Xuwen Qiao, Yuanpeng Zhang, Liting Hou, Xiaoming Yu, Haiwei Cheng, Jin Chen, Qisheng Zheng, Jibo Hou, Guangzhi Tong

## Abstract

Multifunctional CD4^+^ T helper 1 (Th1) cells, producing IFN-γ, TNF-α and IL-2, define a correlate of vaccine-mediated protection against intracellular infection. In our previous study, we found that CVC1302 in oil formulation promoted the differentiation of IFN-γ^+^/TNF-α^+^/IL-2^+^ Th1 cells. In order to extend the application of CVC1302 in oil formulation, this study aimed to elucidate the mechanism of action in improving the Th1 immune response. Considering the signals required for the differentiation of CD4^+^ T cells to Th1 cells, we detected the distribution of innate immune cells and the model antigen OVA-FITC in lymph node (LN), as well as the quantity of cytokines produced by the innate immune cells. The results of these experiments show that, cDC2 and OVA-FITC localized to inter-follicular region (IFR) of the draining lymph nodes, inflammatory monocytes localized to both IFR and T cell zone, which mainly infiltrate from the blood. In this inflammatory niche within LN, CD4^+^ T cells were attracted into IFR by CXCL10, secreting by inflammatory monocytes, then activated by IL-12-secreting cDC2. Above all, CVC1302 in oil formulation, on the one hand, targeted antigen and inflammatory monocyte into the LN IFN in order to attract CD4^+^ T cells, on the other hand, targeted cDC2 to produce IL-12 in order to promote optimal Th1 differentiation. The new finding will provide a blueprint for application of immunopotentiators in optimal formulations.

**Importance:** Along with the development of veterinary immunology, immunopotentiator and delivery system were not simply mixed with antigen, more and more attentions paid on the optimal compatibility in order to induce multifunctional immune responses. As reported, LPS formulated in IFA targeted antigens to the IFR of the LN, and recruited Mo into the IFR, then attracted antigen-specific CD4^+^ T cells, which differentiated into Th1 cells under the IL-12 produced by DC-SIGN^+^ DC(1). Herein, we further found that CVC1302 formulated with Marcol 52 induced enhanced Th1 immunity. Combined with our previous finding that CVC1302 in oil formulation induced improved humoral immunity, we concluded that CVC1302 in oil formulation provided multifunctional immunity, not only higher antibody titers to prevent pathogen infection, but also cellular immunity cytokine to prevent viral shedding.

## Introduction

After infection or vaccination, naïve CD4^+^ T cells in lymph nodes (LN) differentiate into Th1, Th2, Th17, Th9, Th22, or follicular helper T cells (Tfh), classified on the basis of their cytokine and transcription factor profiles(2). Th1 cells play an important role in providing immune protection against a variety of intracellular pathogens(3). Th1 cells secrete IFN-γ, TNF, or IL-2 in various combinations(4). The differentiation of CD4^+^ T cells into Th1 cells requires the recognition of the complex of antigen– MHCII by T cell receptor (TCR) and B7 co-stimulatory molecules by CD28, as well as the cytokine IL-12 via transcription factor STAT4(5–7). The factors mediating T_H_1 responses contain the amount of antigen, the type of antigen-presenting cell (APC) and the cytokine milieu(8).

LNs play a pivotal role in building the innate and adaptive immune response against pathogens(9). The inter-follicular region (IFR) in LNs is the boundary between the B-cell and T-cell zone, and serves as a crossroad that bridges innate and adaptive immunity(10). Oil emulsion has been shown to target antigen to the LN IFR; the toll-like receptor 4 (TLR4) ligand lipopolysaccharide (LPS), formulated in IFA, led to rapid recruitment of inflammatory monocytes to the IFR, which was dependent on CCL2 induced by LPS(1). As reported, CXCR3 ligands, consisting of CXCL9, CXCL10, and CXCL11, play an important role in promoting cell–cell interactions and guiding the intra-LN movements of activated T cells that favor differentiation toward an Th1 cell phenotype(11). Previous studies have reported that LPS-poly(I:C) immunization leads to the induction of CXCL10 in the IFR within 12 hour, mainly produced by inflammatory monocytes, which is dependent on type I IFN derived from LN macrophages(12). This series of the immune reactions creates an inflammatory niche in the IFR leading to the induction of poly-functional Th1 cells. Foot-and-mouth disease virus (FMDV), the agent of foot-and-mouth disease, infects livestock, thereby causing devastating economic losses. CVC1302 in oil emulsion was demonstrated to enhance the FMDV-specific IFN-γ^+^ Th1 immune response(13). We postulated that the mechanisms by which CVC1302 in oil emulsion and LPS in IFA improve the Th1 immune response may be similar. To address this, we elucidated the mechanism of action using a model antigen, OVA or OVA-FITC. Furthermore, we elucidated the contents of the inflammatory niches in the IFR, including the cells, cytokines and chemokines induced by CVC1302 in oil formulation. Our findings provide a new insight for the development of more effective vaccines.

## Materials and Methods

### Mice

Five-week-old, female, pathogen-free BALB/c mice were purchased from Yangzhou University (Yangzhou, China). The study and protocol were approved by the Science and Technology Agency of Jiangsu Province and by the Jiangsu Academy of Agricultural Sciences Experimental Animal Ethics Committee. All efforts were made to minimize animal suffering. All animal studies were performed in strict accordance with the guidelines outlined in the Jiangsu Province Animal Regulations (Government Decree No. 45).

### Antigen, adjuvant, and immunizations

Model antigen OVA and OVA-FITC were both purchased from Solarbio (Beijing, China). CVC1302 is composed of MDP, MPL and β-glucan, all of which were purchased from InvivoGen (San Diego, CA, USA). All components were dissolved in sterile water at the appropriate concentrations to form the aqueous phase, designated CVC1302. OVA or OVA-FITC were diluted with PBS at a concentration of 25 μg/mL. Prior to vaccination, CVC1302 was mixed with OVA or OVA-FITC at a ratio of 1:2, then emulsified with Marcol 52 mineral oil at a ratio of 1:2; both of these were designated OVA-CVC1302-M52. OVA or OVA-FITC mixed directly with Marcol 52 mineral oil at a ratio of 1:2 were designated OVA-M52. OVA or OVA-FITC mixed with CVC1302 at a ratio of 1:2, with PBS then added to 100 μL, were designated OVA-CVC1302. OVA or OVA-FITC (2 μL) diluted into PBS to 100 μL, were designated OVA.

Mice were immunized intramuscularly in the quadriceps muscles of each hind leg with 50 μg OVA (OVA; grade V, Sigma) with or without 1 μL CVC1302 in 100 μL PBS or emulsified 1:2 in Marcol 52 mineral oil.

### LN re-stimulation

1 × 10^6^ total cells from draining LNs were cultured at 37°C in 5% CO_2_ for 3 days in 96-well round bottom plates containing 100 μg/mL OVA protein in 200 μL of complete RPMI (containing 10% heat-inactivated fetal bovine serum (FBS), 100 U/mL penicillin, 100 μg/mL streptomycin), and supernatants were collected and stored at −80°C. To assess polyfunctional T cell responses, 1 × 10^6^ total cells from draining LNs were cultured at 37°C in 5% CO_2_ for 72 hours in 96-well round bottom plates containing 100 μg/mL OVA protein in 200 μL of complete RPMI, then incubated with 2 μg/mL protein transport inhibitor cocktail (500×) (eBioscience) in complete RPMI at 37°C in 5% CO_2_ for 4 hours.

### Enzyme-linked immunosorbent assay (ELISA)

Culture supernatants from restimulated LNs were harvested and the presence of IFN-γ was tested with commercial mouse IFN-γ immunoassay ELISA kits (Booster Biological Technology, LTD., Wuhan, China) in accordance with the manufacturer’s protocols. The concentrations of IFN-γ in the samples were determined based on the standard curves.

For OVA-specific ELISA, sera were harvested from immunized mice at 28 days post-immunization (dpi). ELISA plates were coated with 10 μg/mL OVA at 4°C overnight, then blocked with 5% skimmed milk at 37°C for 2 hours. Duplicate sera samples at a 1:200 dilution were incubated at 37°C for 1 hour, followed by incubation with goat anti-mouse IgG1- or IgG2a-adsorbed HRP (Southern Biotech) for 1 hour. Then, 3, 3′, 5, 5″-tetramethylbenzidine (TMB) substrate solution (Biopanda Diagnostics, UK) was added at 37°C for 15 min, and the reaction was stopped with 2 M H_2_SO_4_. The OD450 was determined using an ELISA reader (BioTek, USA).

### Flow cytometry

Draining LNs were harvested and cut into small fragments. Single lymphocytes were prepared using a Medimachine system with a Medicon (50 μm) disaggregator (BD, USA). Cells were washed twice in ice cold PBS (containing 2% FBS), blocked with anti-CD16/32 at 4°C for 10 min, and stained with the indicated antibodies at 4°C for 30 min. For intracellular cytokine staining, after surface staining, cells were fixed and permeabilized with intracellular fixation and permeabilization buffer (ThermoFisher Scientific, USA) and then stained for intracellular cytokines. Cells were washed and resuspended in PBS (containing 2% FBS). Cells were analyzed with a BD Accuri C6 instrument (BD Biosciences, USA). Data analyses were performed using FlowJo version 10 software. The antibodies are listed in Supplemental Table 1.

### Confocal microscopy

LNs were collected at the appropriated time points and fixed in periodate-lysine-paraformaldehyde (PLP) at 4°C for 4 hours, then embedded in optimum cutting temperature compound (OCT). Cryosections (6 μm) of LNs were obtained using a CM1950 cryostat (Leica Microsystems, Germany). Tissue samples were captured on Superfrost plus microscope slides (ThermoFisher Scientific, USA), then washed with PBS three times to remove any residual OCT. The cryosections were fixed using PBS plus 3% formaldehyde for 10 min at room temperature, washed twice with PBS, and permeabilized with PBS plus 3% BSA/1% saponin (permeabilization buffer) for 30 min at room temperature. Tissue sections were then stained with the indicated antibodies for 1 hour at room temperature in the dark. In some circumstances, samples were incubated with secondary antibodies for another 1 hour at room temperature in the dark. Sections were visualized and photographed with a Zeiss LSM700 confocal microscope (objective, 320) at room temperature, and images were acquired with Zeiss LSM image browser software (Zeiss).

### Real-time quantitative PCR (RT-qPCR)

RNA was extracted from LNs and used for RT-qPCR using a Prime Script™ II Strand cDNA Synthesis kit (Takara, Dalian, China), in accordance with the manufacturer’s protocol. Real-time qPCR was performed for each cDNA sample in duplicate using Bright Green 2 × qPCR Master Mix in a Roche Light Cycler®480. The thermal cycling program was as follows: an initial denaturation step at 95 °C for 10 min, followed by 40 cycles of 15 s at 95 °C and 1 min at 60 °C. The expression level of GAPDH was used as the internal control, and gene expression was measured by the relative quantity as described previously. The primers for all of the detected genes are listed in Supplemental Table 2.

### Statistical Analysis

Statistical analysis was performed using GraphPad Prismversion5 (GraphPad Software, San Diego, CA, USA). Differences among groups were assessed using one-way analysis of variance followed by Tukey’s post-hoc t-test. Differences between groups were assessed using a Student’s t-test. Values of *p* < 0.05 were considered statistically significant. All data shown in the manuscript are expressed as the means ± standard error of the mean (SEM).

## Results

### CVC1302 in oil formulation leads to a more robust Th1 response

To identify whether CVC1302 alone, without emulsification, could induce a significant Th1 response against intracellular infections, we immunized mice intramuscularly in the quadricep muscle of each hind leg with 50 μg OVA (OVA; grade V, Sigma), with or without 1 μL CVC1302, in 100 μL PBS or emulsified 1:2 in Marcol 52 mineral oil. At 7 dpi, a significantly higher concentration of IFN-γ was observed from draining LN cells cultured with OVA for 3 days, only in mice immunized with OVA-CVC1302-M52, not in groups of mice immunized with OVA alone, OVA-CVC1302, or OVA-M52 (Figure 1A).

**Fig. 1.**
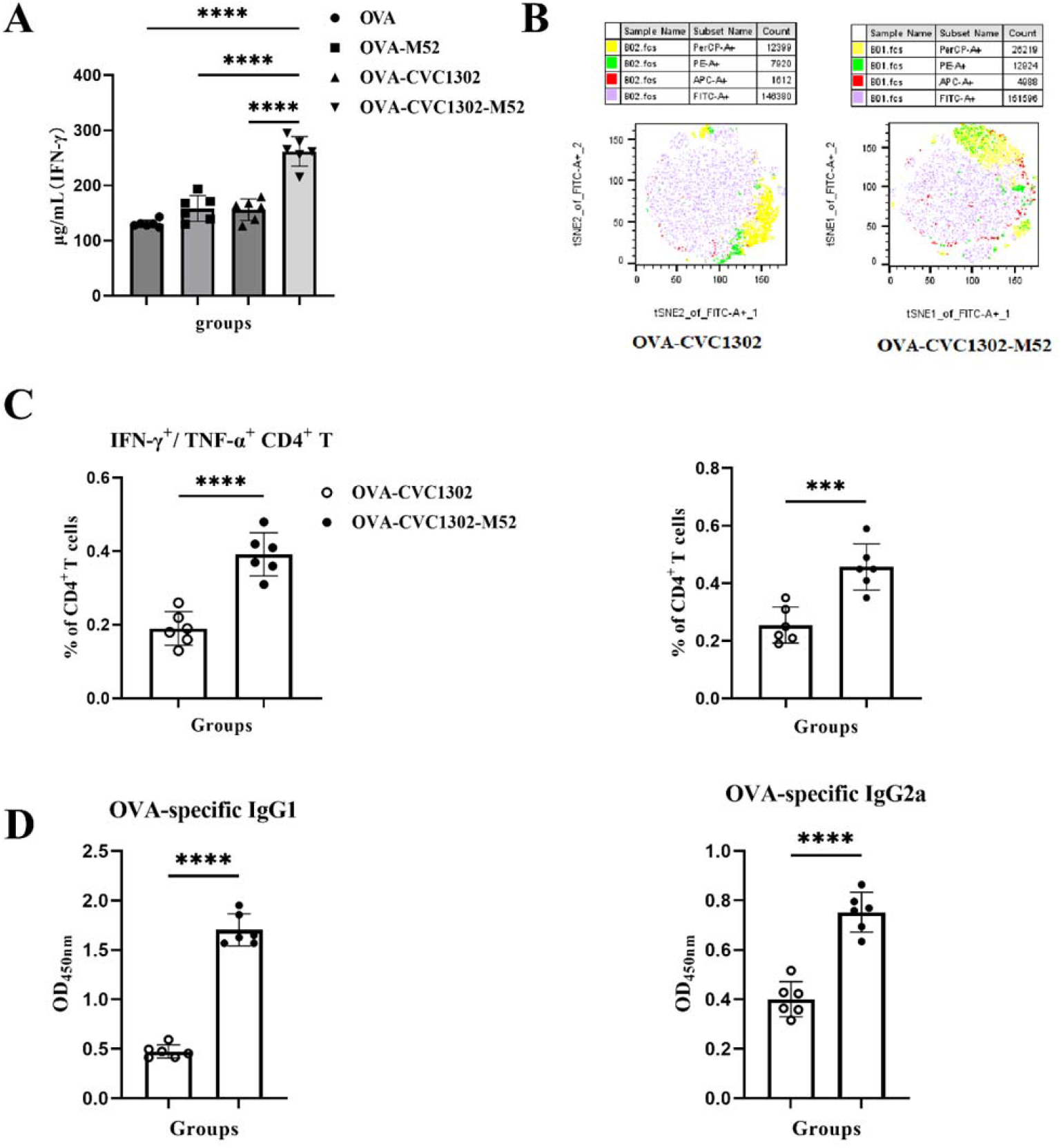
CVC1302 in oil formulation leads to a more robust Th1 response. BALB/c mice were immunized intramuscularly in the quadriceps muscles with 50 μg OVA with or without 1 μL CVC1302 in PBS or emulsified 1:2 in Marcol 52, and LNs were harvested at 6 dpi. (A) LN cells were cultured with OVA, and IFN-γ in the supernatant was measured by ELISA. (B) LNs cultured with OVA as in (A) were re-stimulated with phorbol 12-myristate 13-acetate (PMA) and ionomycin and stained for intracellular cytokine. tSNE analysis of IFN-γ, TNF-α, and IL-2 -producing CD4^+^T cells. (C) Percentages of IFN-γ^+^/ TNF-α^+^ -producing CD4^+^ T cells and IFN-γ^+^/IL-2^+^ -producing CD4^+^ T cells in mice immunized with OVA-CVC1302 or OVA-CVC1302-M52. (D) Relative OVA-specific immunoglobulin G (IgG1 and IgG2a) levels were measured by ELISA at 42 dpi. Data are presented as mean ± SEM; \*\*\**P*≤ 0.001, \*\*\*\**P*≤ 0.0001.

It is well established that differences in the combination of cytokines (IFN-γ, TNF, and IL-2) produced by individual Th1 cells have profound implications for their capacity to mediate effector function and their capacity to be sustained as memory, or both(14). Considering that IFN-γ and TNF synergize in their capacity to mediate the killing of pathogens, and IL-2 strongly enhances the expansion of CD4^+^ and CD8^+^ T cells leading to a more efficient effector response, the frequency of multifunctional Th1 cells simultaneously producing IFN-γ, TNF, and IL-2 correlated most closely with protection induced by vaccines(14–16).

Using multiparameter flow cytometry, the frequencies of distinct Th1 cells based on the combination of cytokines (IFN-γ, TNF, and IL-2) were analyzed in OVA-cultured lymphocyte cells after immunization with OVA-CVC1302-M52, OVA alone, OVA-CVC1302 or OVA-M52. Significantly higher percentages of IFN-γ^+^ TNF^+^ and IFN-γ^+^ IL-2^+^ Th1 cells were observed in mice immunized with OVA-CVC1302-M52 than those immunized with OVA alone, OVA-CVC1302 or OVA-M52 (Figure 1B and 1C).

As known that IFN-γ promote effector B cells secreting Ig2a, so we detected the titers of isotype antibody (IgG1 and IgG2a). From the results of Figure 1D, CVC1302 formulated with oil formulation induced higher titers of IgG1 and IgG2a, which keep in line with our previous study.

### Oil formulation targets antigen to the IFR of LNs

Because CVC1302, containing pattern recognition receptors (PRRs) agonists, adjuvanted with OVA, did not induce significantly enhanced Th1 responses in mice when compared with mice immunized with OVA alone, we focused on the distribution of antigen in LNs induced by Marcol 52 mineral, as it was demonstrated that IFA, as an oil formulation, could promote antigen targeting to the IFR of LNs. To investigate this, we used OVA-FITC as antigen and immunized mice with CVC1302 formulated with or without Marcol 52 mineral, then observed the distribution of OVA-FITC in the LN by confocal microscopy.

From the results, we found that OVA-FITC was mainly distributed in the IFR of the LN in mice immunized with OVA-CVC1302-M52 at 1 dpi. By contrast, OVA-FITC was mainly in T cells of the LN in mice immunized with OVA alone or OVA-CVC1302, which demonstrated that the change in the pattern of antigen distribution was based on the oil formulation not CVC1302 (Figure 2).

**Fig. 2.**
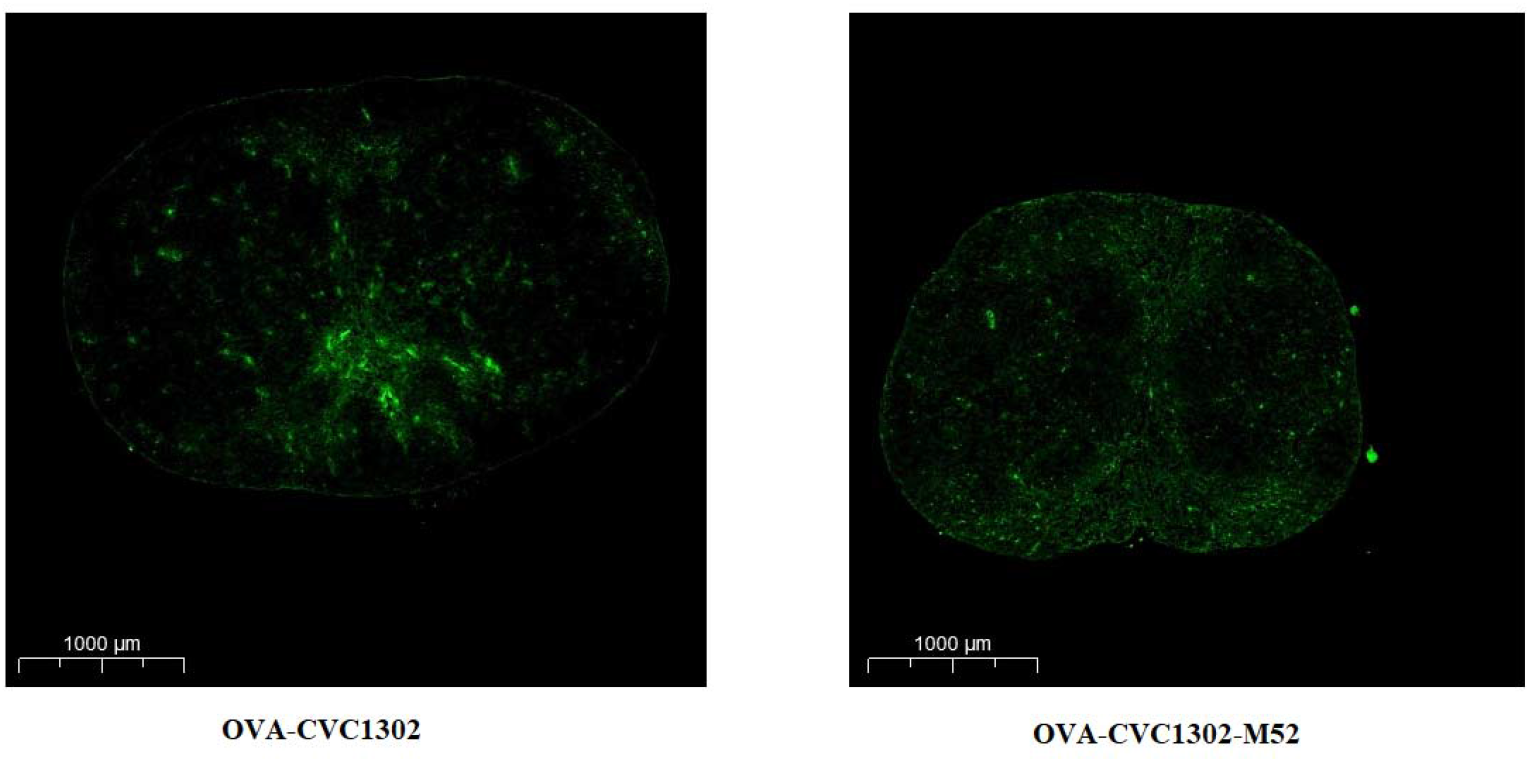
Oil formulation targets antigen to the IFR of LNs. BABL/c mice were immunized with OVA-FITC and CVC1302 in PBS (OVA-CVC1302, left) or OVA-FITC and CVC1302 emulsified in Marcol 52 (OVA-CVC1302-M52, right). LNs were harvested after 24 h, and analyzed by confocal microscopy.

It was established that the enhanced immune responses induced by antigen formulated with oil formulation were mainly due to the injection depot resulting in sustained antigen release. However, IFA, as an oil formulation, did not influence the number of OVA^+^ cells in the LNs at the early period after immunization, aside from the sustained release of antigen. We therefore questioned whether Marcol 52 mineral, as an oil formulation, has an identical mechanism of action to IFA. We determined the total number of OVA^+^ cells and the number of OVA^+^ APCs. In line with our assumption, no significant difference in the number of OVA^+^ cells at 1, 2, and 3 dpi, which means that the difference in the Th1 response at 7 dpi was not due to the amount of antigen, and may instead be due to the pattern of antigen distribution (Figure 3A).

**Fig. 3.**
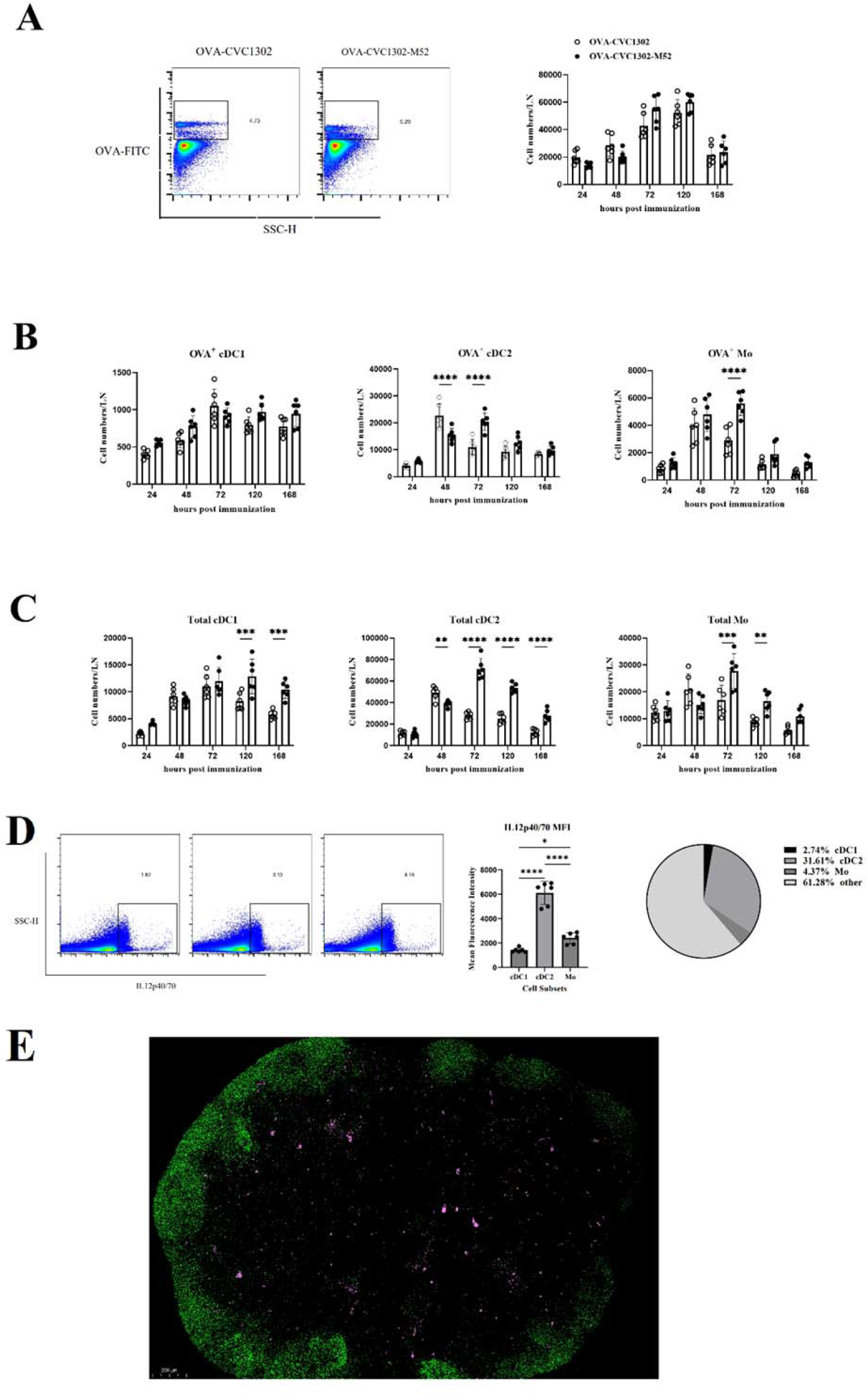
cDC2 are the dominant cell type responsible for the robust Th1 immune response induced by CVC1302 in oil formulation. (A, B and C) BABL/c mice were immunized with OVA-FITC-CVC1302 or OVA-FITC-CVC1302-M52. LNs were harvested at the indicated time points and analyzed by flow cytometry. (A) (left) Representative flow cytometry plot showing total OVA^+^ cells 24 h post-immunization gated on FSC/SSC single cells. (right) OVA^+^ cells numbers in LNs at the indicated time-points post-immunization. (B) OVA^+^ cell numbers of the indicated subsets. (C) Total cell numbers of the subsets from (B). (D) LNs were harvested 72 h post-immunization. Shown are representative flow cytometry plots of IL12p40/70^+^ cells (left), MFI of IL12p40/70^+^ cells of the indicated subsets from (B) (middle), and percentages of IL12p40/70^+^ cells of the indicated subsets from (B) (right). (E) LNs from BABL/c mice 72 h post-OVA-CVC1302-M52 immunization were stained B220 (green) and IL12p40/70 (pink) and analyzed by confocal microscopy. Data are presented as mean ± SEM; \*\**P*≤ 0.01, \*\*\**P*≤ 0.001, \*\*\*\**P*≤ 0.0001.

### Conventional dendritic cells 2 (cDC2) is responsible for the robust Th1 response induced by CVC1302 in an oil formulation

Regarding the potential effect of the oil formulation on the antigen distribution pattern, we next investigated whether the oil formulation may influence APCs localization induced by CVC1302. To achieve this, we immunized mice with OVA-CVC1302 or OVA-CVC1302-M52, then isolated LNs at 1 dpi and visualized the distribution patterns of APCs using confocal microscopy. We observed that compared with the naïve group, cDC1 re-localized to the deep T zone, cDC2 mainly repositioned to the IFR and B zone, and large numbers of monocyte (Mo) infiltrated and distributed within both the IFR and the T zone of the LNs, after either OVA-CVC1302 or OVA-CVC1302-M52 immunization (Figure 4).

**Fig. 4.**
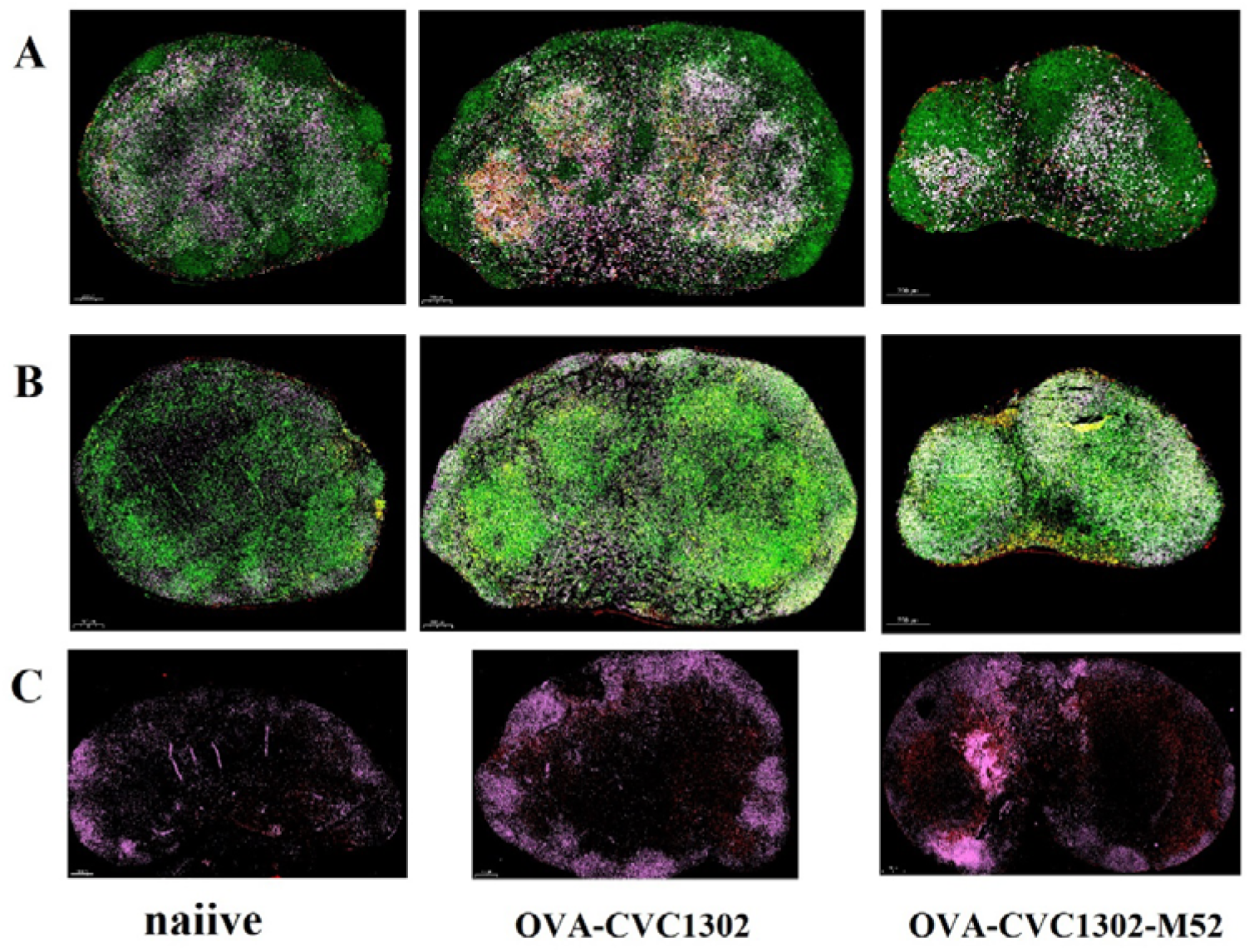
cDC re-localization and Mo influx during CVC1302 in oil formulation immunization. BABL/c mice were immunized intramuscularly in the quadriceps muscles with OVA-CVC1302 or OVA-CVC1302-M52 and LNs were analyzed 24 h post-immunization by confocal microscopy. (A) LNs were stained with CD11c (green), CD103 (red) and CD8α (pink). Migrated cDC1 (yellow) and resident cDC1 (white) were shown in the merged picture. (B) LNs were stained with CD11c (green), CD11b (red) and CD172α (pink). cDC2 (white) were shown in the merged picture. (C) LNs were stained with B220 (pink) and CCR2 (red). Scale bars denote 200 μm.

Considering the distribution of antigen and APCs (cDC2 and Mo), we hypothesized that cDC2 was the main APC responsible for robust Th1 responses. To confirm this, we analyzed the recruitment levels of OVA^+^ APCs. As shown in Figure 3B, we found that at the earlier time points of 24 h and 48 h post-immunization, Mo increased more rapidly in mice immunized with OVA-CVC1302 than in mice immunized with OVA-CVC1302-M52, but at the later time points of 48 h and 72 h post-immunization, there was a rapid increase in Mo in mice immunized with OVA-CVC1302-M52, compared with mice immunized with OVA-CVC1302. By comparison, there was no significant difference detected in the amounts of OVA^+^ cDC1 between the groups immunized with OVA-CVC1302 and OVA-CVC130-M52. At the sampled time points of 24 h, 48 h, and 72 h post-immunization, cDC2 increased more rapidly in the mice immunized with OVA-CVC1302-M52 than in those immunized with OVA-CVC1302. Even total Mo was quickly infiltrated into the LNs; however, compared with cDC2, the numbers of OVA^+^ Mo were much lower than those of OVA^+^ cDC2 in both mice immunized with OVA-CVC1302 and OVA-CVC1302-M52.

It is known that the induction of a Th1 response requires the collaboration of antigen, APCs, and the innate cytokine milieu(17). Considering that IL-12 predominantly induces the differentiation of naïve CD4^+^ T cells into Th1 cells, the key IL-12-producing cell population was determined using flow cytometry. At 120 h post-immunization, we found that a higher percentage of IL-12-producing cDC2 in the mice immunized with OVA-CVC1302-M52 than in those immunized with OVA-CVC1302. Mo also produced IL-12, but cDC2 and Mo showed a significant difference in their percentage abundance and mean fluorescent intensity (MFI) (Figure 5).

**Fig. 5.**
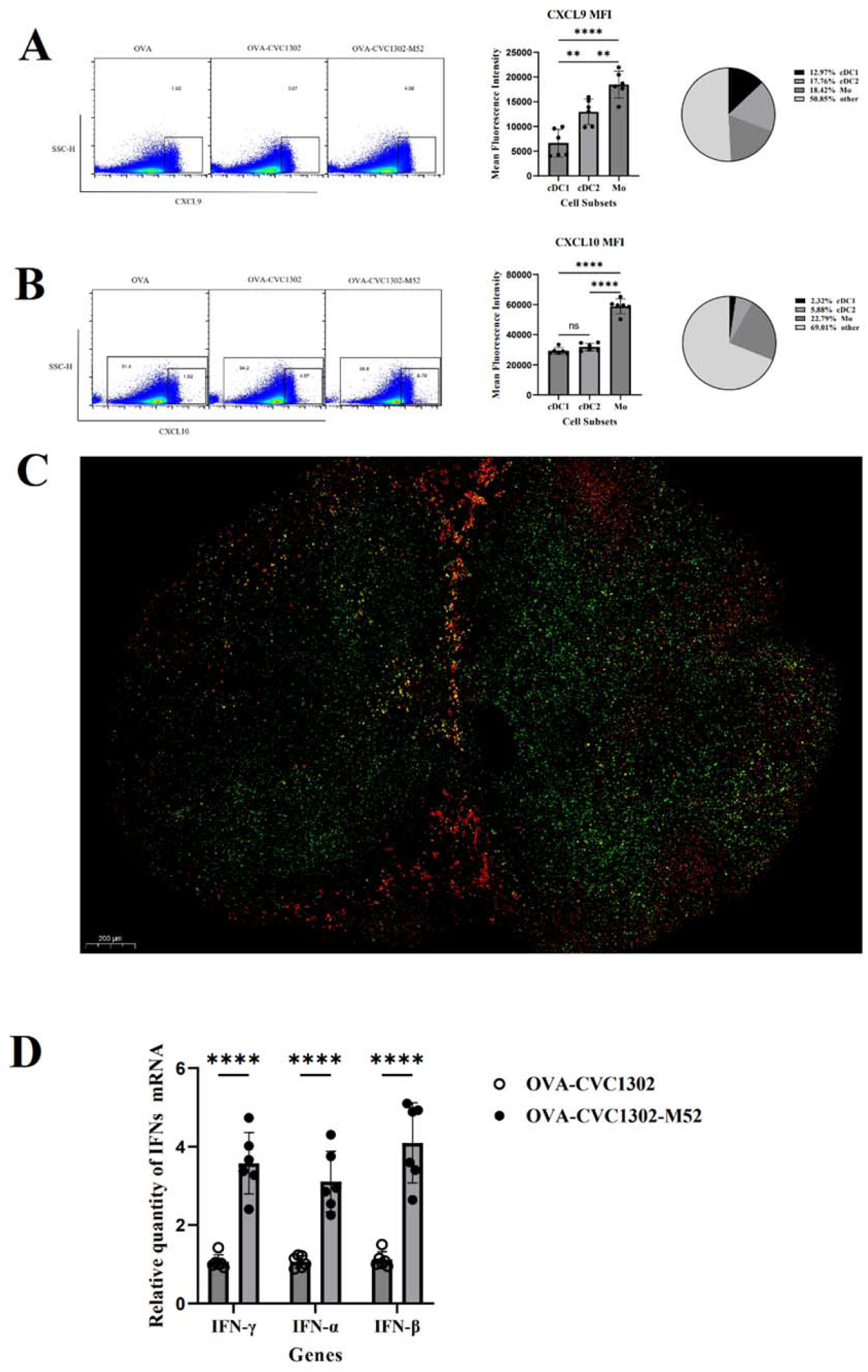
Mo was the prominent source of CXCL9 and CXCL10 dependent on IFN-γ and type I IFN. BABL/c mice were immunized intramuscularly in the quadriceps muscles with OVA-CVC1302 or OVA-CVC1302-M52 and LNs were analyzed 72 h post-immunization by flow cytometry and confocal microscopy. (A) Representative plot of CXCL9^+^ cells 72 h post-immunization (left). MFI of CXCL9 expression with the indicated cell subset (middle). Percentage of CXCL9 expression with the indicated cell subset (right). (B) Representative plot of CXCL10^+^ cells 72 h post-immunization (left). MFI of CXCL10 expression with the indicated cell subset (middle). Percentage of CXCL10 expression with the indicated cell subset (right). (C) LNs from BABL/c mice 72 h post-OVA-CVC1302-M52 immunization were stained CXCL9 (red) and CXCL10 (green) and analyzed by confocal microscopy. (D) The expression levels of IFN-γ and type I IFN were detected by real-time RT-PCR. LNs were isolated at 48 h post-immunization, and RNA was extracted from these cells by using Trizol. Data are presented as mean ± SEM; \*\**P*≤ 0.01, \*\*\*\**P*≤ 0.0001.

Based on the results of confocal microscopy and flow cytometry, we concluded that cDC2 was the dominant cell type responsible for the robust Th1 immune response induced by CVC1302 in oil formulation.

### Mo promotes antigen-specific CD4^+^ T cells localization into the IFR of LNs

It was revealed that the expression pattern of two Th1 cell-associated chemokine receptor CXCR3 ligands, CXCL9 and CXCL10, guides the intra-LN movements of activated T cells that favors differentiation toward an IFN-γ-producing Th1 cell phenotype(12). Considering the distribution of Th1 cells, we predicted that one or both subsets of CXCL9- and CXCL10-producing cells might be localized in the IFR, thereby attracting CD4^+^ T cells into the IFR to be activated by OVA^+^ cDC2 that promotes Th1 differentiation.

Previous studies have found that mice immunized with soluble antigen, TLR ligands, DC-bearing antigen lead to the induction of CXCL10 in the IFR, the medullary and sub-capsular sinus regions, and CXCL9 in the IFR and some T zone areas at 36 h post-immunization. Considering that Mo and cDC2 localized in the IFR after immunization with CVC1302 in oil formulation immunization, we detected the ability of Mo and cDC2 in producing CXCL9 and CXCL10 by flow cytometry at 72 h post-immunization. As shown in Figure 5A and B, both Mo and cDC2 have the ability to produce CXCL9 and CXCL10, but compared in the MFI, there was a significant difference among Mo and cDC2. From the results of confocal microscopy (Figure 5C), we found that CXCL9 and CXCL10 producing cells mainly distributed in IFR and T zone, so we concluded that CXCL9 and CXCL10 were the dominant chemokines responsible for attracting antigen-specific CD4^+^ T cells into the IFR of LNs.

From the results, we concluded that infiltrated Mo was the dominant cell type responsible for the localization of antigen-specific CD4^+^ T cells into the IFR of LNs through producing CXCL9 and CXCL10.

It has been well established that CXCL9 and CXCL10 production is regulated by IFN-γ and type I IFN, respectively. At 48 h post-immunization, LNs were sampled from mice immunized with OVA-CVC1302-M52 and OVA-CVC1302, then the relative quantity of type I IFN mRNAs were measured using qPCR. As shown in Figure 5D, the IFN-γ and type I IFN genes in OVA-CVC1302-M52-immunized mice had higher transcription levels as compared with those in OVA-CVC1302 mice.

Since we observed that the dominant CXCL9- and CXCL10-producing Mo localized in IFR of the LNs in responsible to OVA-CVC1302-M52 immunization, we wonder whether the antigen-specific CD4^+^ T cells would be attracted into IFR, then received MHCII-antigen transferred by cDC2 and IL-12 produced also by cDC2 that promote Th1 differentiation. In order to visualize antigen-specific CD4^+^ T cell localization in mice immunized with OVA-CVC1302-M52, LNs were harvested at 7 dpi and stained with indicated fluorescent antibody in Figure 6. From the results, we found that OVA-specific CD4^+^ T cells in OVA-CVC1302-M52 immunized mice were distributed in IFR and T zone in OVA-CVC1302-M52 immunized mice, and OVA-specific CD4^+^T cells in OVA-CVC1302 immunized mice were distributed only in T zone in OVA-CVC1302-M52 immunized mice.

**Fig. 6.**
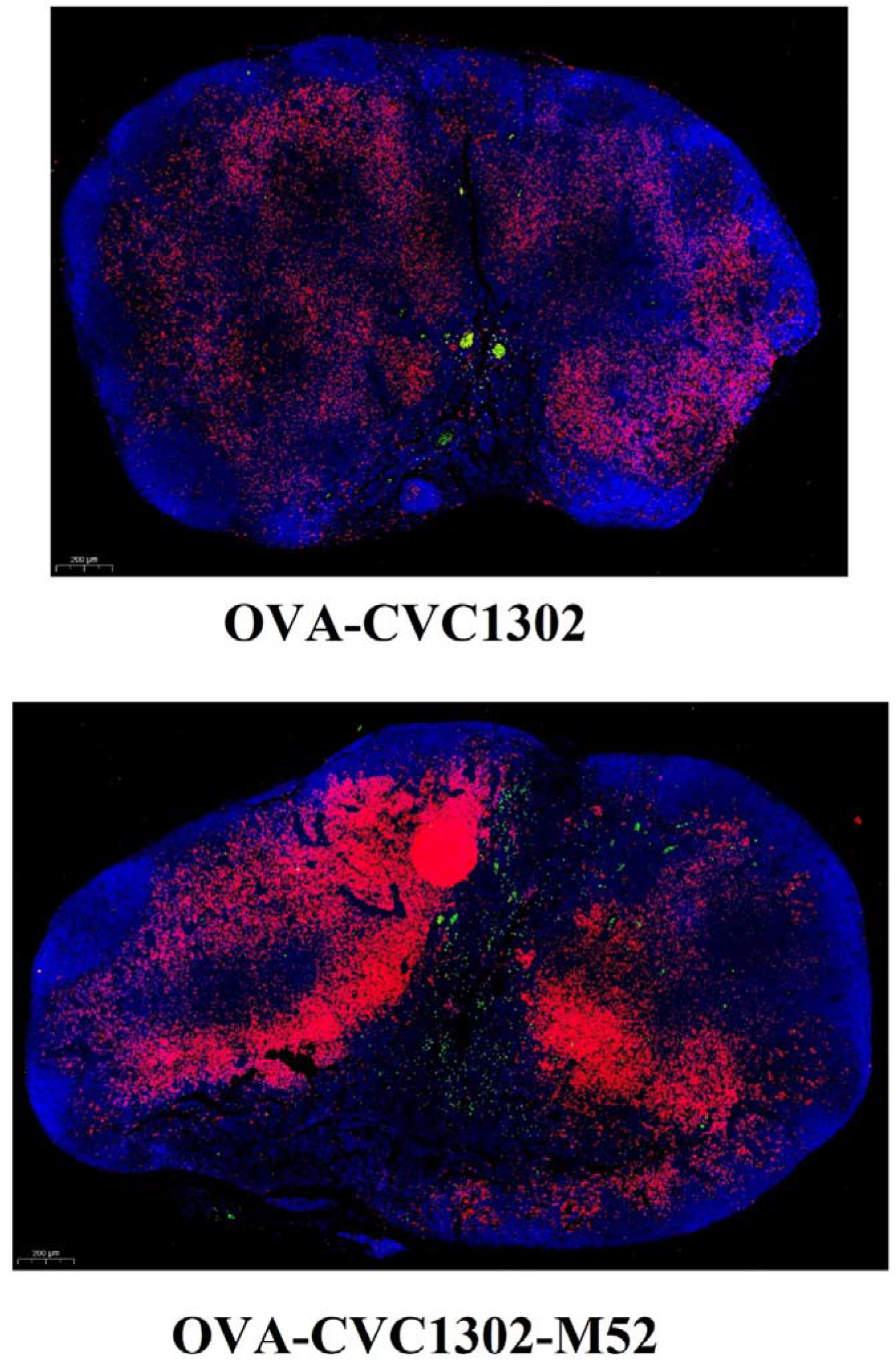
CVC1302 in oil formulation promotes antigen-specific CD4^+^ T cells localization into the IFR. BABL/c mice were immunized with OVA-CVC1302 and OVA-CVC1302-M52 and LNs were harvested 7 dpi and stained with OVA (329-337)-FITC (green), CD4 (red), and DAPI (blue). OVA-specific CD4^+^ T cells (yellow) were shown in the merged picture.

## Discussion

In a previous study, it was demonstrated that LPS in IFA, an oil formulation, promoted antigen targeting in the LN IFR, where CD4^+^ T cells, attracted by CXCL10 produced by infiltrated Mo, differentiated into Th1 cells under the activation of antigen mainly presented by DC-SIGN^+^ DCs and IL-12 also by DC-SIGN^+^ DCs. It is known that Th1 cells are critical for immunity against intercellular pathogens, potentially preventing viral shedding. In our previous study, we found that CVC1302 could promoted long-term humoral immunity induced by FMDV inactivated vaccine in order to protect immunized pigs against FMDV(13). However, advances in immunology have revealed that ideal vaccines should provide not only humoral immunity to against pathogens, but also cellular immunity to prevent viral shedding. Considering that CVC1302, a complex of PRRs agonist available for clinical use, is formulated with Marcol 52 mineral oil and this oil formulation is thought to induce a robust Th1 immune response, so we utilized OVA as a model antigen to verify our hypothesis.

We found that Marcol 52, as a water-in-oil formulation, promoted OVA targeting to the IFR of LN and CVC1302, as a complex of PRRs agonist, attracted Mo to infiltrate the IFR and T zone and cDC2 to distribute to the IFR and B zone. Utilizing flow cytometry, we discovered that Mo produced CXCL10 and cDC2 produced IL-12. Under the innate immune microenvironment generated by CVC1302 in oil formulation, firstly, cDC2 was activated by CVC1302 then captured, degraded, and presented OVA via MHCII to naïve T cells, which translocated into the IFR attracted by CXCL10. Second, activated T cells differentiated into Th1 cells in response to IL12 secreted by cDC2. Finally, Th1-cell mediated cellular immunity protected against intracellular pathogens through secreted cytokines such as IFN-γ, IL-2, and TNF.

Previous work has established that induction of the CXCR3 ligand CXCL9 and CXCL10 was dependent on IFN-γ and type I IFN, respectively(18–20). Higher relative quantities of IFN-γ, IFN-α and IFN-β mRNA were detected in mice immunized with OVA-CVC1302-M52 than in those immunized with OVA-CVC1302. Furthermore, in our previous study, it was demonstrated that CVC1302 downregulated the expression of Nmi, which was responsible for negative regulation of type I IFN(21). Using flow cytometry, we confirmed that CXCL10 predominatly originated from Mo. Even the percentage of CXCL9 producing Mo and cDC2 had no significant difference in mice immunized with OVA-CVC1302-M52, the MFI of CXCL9 from Mo and cDC2 had a significant difference, so we concluded that Mo was the dominant cells that attracted naïve T cells to translocate to the IFR of the LNs. Because antigen and the innate cytokine IL-12 are known to induce Th1 responses(22), we investigated the original cell subset of IL-12 by determining the percentages and MFIs of IL-12p70^+^ cDC1, cDC2, and Mo. From the results, we found that IL-12p70 predominantly originated from cDC2 but that Mo and cDC1 also poduced IL-12p70(23).

An ideal vaccine should simultaneously provide both humoral and cellular immunity to effectively mimic the immune response of a natural infection. Humoral immunity is mediated by B cells, and cellular immunity relies on Th1 cells. In our previous study, we discovered that CVC1302 induced long-term humoral immunity through the positive selection of enhanced Tfh, promoting germinal center B cells to differentiate into high-affinity long-lived plasma cells, prolonging the lifetime of antigen-specific long-lived plasma cells. CVC1302 mediates cDC1 cross-presentation to promote CD8^+^ T cell immunity (unpublished). This study further demonstrated that CVC1302 in oil formulation could improve Th1 immunity. Overall, we illustrated the immunity mechanism of CVC1302 in promoting multifunctional immune responses, which provides new insights into the development of novel immunopotentiators in optimal formulations.

## Author Contributions

LD, QS, JC and JH designed the experiment. Sampling of LNs and preparation of lymphocytes were mainly performed by LD, LT and XY. LD, QX, PY and HC analyzed the results with guidance from QS and JH and wrote the main manuscript text. All authors took part in discussion and interpretation of results. All authors read, advised and approved the final manuscript.

## Acknowledgements

This work was supported by the National Natural Sciences Foundation of China (32102690), Jiangsu Agricultural Science and Technology Innovation Fund (CX (21)3135).

